# Pulmonary Vascular Endothelial Dysfunction is Induced by Non-Pulsatile Pulmonary Blood Flow in an Ovine Classic Glenn Model

**DOI:** 10.1101/2025.10.08.681256

**Authors:** Jonathan Hyde, Michael A. Smith, Naveen Swami, John H. Hwang, Yenchun Chao, Jason Boehme, Gary W. Raff, Casper Noah Nilsson, Wenhui Gong, Gail H. Deutsch, Eric G. Johnson, Ting Wang, Stephen M. Black, Sanjeev A. Datar, Emin Maltepe, Jeffrey R. Fineman

## Abstract

**Background:** Pulmonary vascular disease (PVD) in patients with single ventricular heart disease following the partial cavalpulmonary connection (Glenn) is a significant source of morbidity. However, the etiology of pulmonary vascular endothelial cell (EC) dysfunction, an established precursor to PVD, is incompletely understood but may involve abnormal blood flow patterns, hypoxemia, and polycythemia.

**Hypothesis:** Utilizing an ovine Glenn model, we hypothesized that non-pulsatile pulmonary blood flow (PBF) induces pulmonary vascular EC dysfunction, independent of hypoxemia or polycythemia.

**Methods:** Seven lambs (6-8 weeks old) underwent a Glenn procedure. Eight weeks later, Glenn and age-matched controls were studied. The response to the endothelium-dependent vasodilator acetylcholine (Ach) was determined in isolated pulmonary arteries (PA). Nitric oxide (NO) and endothelin-1 (ET-1) signaling was determined in right lung tissues. Indices of cell proliferation, angiogenesis, and apoptosis were determined in PA endothelial cells (PAECs). Comparisons were made by unpaired t-test and ANOVA.

**Results:** There were no differences in age, hemoglobin, or oxygen saturation between groups. Mean PA pressure and left PA flow were higher, and right lung blood flow was lower in Glenn lambs compared to controls (p<0.05). All other baseline hemodynamics were similar. Glenn PAs had impaired relaxation to Ach. Glenn lung NO metabolite levels (NOx) and eNOS protein were lower, and ET-1 levels and prepro-ET-1 protein were higher than controls (p<0.05). Glenn PAECs had higher rates of proliferation and angiogenesis, and decreased apoptosis (p < 0.05).

**Conclusions:** The initiation of non-pulsatile PBF following the Glenn induces early EC dysfunction independent of hypoxemia and polycythemia.

## Introduction

Despite improvements in the management of single ventricle heart disease (SVHD), lifespan remains curtailed, and patients continue to suffer significant morbidities^1–8^. Fundamental to these morbidities and early mortality is the development of vascular dysfunction^1,3–5,7,8^. Although the mechanisms underlying this vascular dysfunction are not fully understood, they are likely influenced in part by the staged surgical palliation these patients receive. The anatomy of SVHD is quite heterogenous, and thus the details of the initial surgical intervention will vary. However, in the overwhelming number of cases, an intervention in the neonatal period is required to establish adequate sources of pulmonary and systemic blood flows, and ensure unobstructed, adequate intracardiac mixing of blood. Within the first year of life (usually 3-6 months of age), the infant undergoes a second procedure, the partial cavalpulmonary connection (Glenn procedure), that begins the process of separating the pulmonary and systemic circulations. In the Glenn, the superior vena cava (SVC) is connected directly to the pulmonary artery, bypassing the heart^9^. Thus, the head and upper extremity venous blood return from the SVC becomes the sole, non-pulsatile source of pulmonary blood flow. The goal of the Glenn procedure is to decrease the volume load of the single ventricle, and thereby decrease its workload, until the child eventually undergoes the total cavalpulmonary connection (Fontan procedure). In the Fontan circulation, the inferior vena cava (IVC) blood flow is directed to the pulmonary arteries, completing the separation of systemic and pulmonary circulations. This further decreases the workload of the single ventricle and removes intracardiac mixing and the associated hypoxemia^10,11^. At this point the non-pulsatile pulmonary blood flow represents a full cardiac output^12^ .

The period between the establishment of the partial cavalpulmonary connection (Glenn) and the total cavalpulmonary connection (Fontan) is critical for SVHD patients and often marked by the development of pulmonary vascular disturbances that contribute to lifelong SVHD morbidities and shortened lifespan^13–15^. For example, pulmonary vascular disease in patients with SVHD following the Glenn procedure may result in upper body swelling, headaches, lymphatic morbidities, hypoxemia, decreased cardiac output, and limited palliative options, including the inability to proceed to the Fontan procedure^16–18^ . Pulmonary vascular endothelial cell (EC) dysfunction, often characterized by decreased bioavailable nitric oxide (NO), increased endothelin-1 (ET-1), and an EC hyperproliferative, angiogenic, and anti-apoptotic phenotype, is an established precursor to pulmonary vascular disease^19^. Pulmonary vascular EC dysfunction is well characterized following the Fontan. In fact, some studies suggest that the degree of pulmonary vascular EC dysfunction correlates with poor outcomes^20–23^. However, data are quite limited following the Glenn procedure, prior to the Fontan^23–25^. In addition, the role of potential drivers of this pathology, which include hypoxemia, polycythemia, inflammation, and aberrant pulmonary blood flow patterns, are unclear.

It is increasingly appreciated that “normal” physiologic blood flow patterns are required to maintain vascular homeostasis^26,27^. Conversely, perturbed flow patterns, including those that result from congenital heart disease (CHD), expose ECs to abnormal mechanical forces that drive pulmonary vascular endothelial dysfunction^28–30^. Unfortunately, progress towards mechanistic insights into SVHD pathologies and novel treatment advances has been hampered by the dearth of clinically relevant animal models^31^. Thus, the objective for this study was to utilize a novel large animal model of SVHD, the “classic” Glenn lamb, in which non-pulsatile blood flow is established to the right lung only, to characterize aberrant pulmonary vascular endothelial function in the setting of non-pulsatile pulmonary blood flow. To this end, we utilized an integrated *in-vivo* (whole animal hemodynamics and imaging), *ex-vivo* (isolated pulmonary artery reactivity), and *in-vitro* (primary pulmonary artery endothelial cell, PAEC, culture) investigative platform to comprehensively study the hypothesis that non-pulsatile pulmonary blood flow, independent of hypoxemia and polycythemia, will induce significant alterations in endothelial function, including a decrease in NO signaling and an increase in ET-1 signaling, that ultimately drive pulmonary vascular dysfunction in patients with the Glenn circulation.

## Materials & Methods

### Surgical Preparation, Animal Care, and Hemodynamics

A total of seven mixed-breed Western lambs (6-8 weeks old) were anesthetized with midazolam and ketamine, intubated and mechanically ventilated. Anesthesia was maintained with isoflurane. Using aseptic technique, a median sternotomy, partial thymectomy, and pericardiotomy were performed. The superior vena cava and right pulmonary artery were identified and dissected free from their attachments. The azygous vein was divided. Intravenous heparin was administered (300 U/kg), and 16F to 20F venous cannulas were placed to bypass the superior vena cava to the right atrium. The right pulmonary artery was divided near the pulmonary bifurcation and the proximal end was oversewn. The superior vena cava was similarly divided at the cavoatrial junction, and the atrial end was oversewn. The superior vena cava was then anastomosed to the right pulmonary artery in an end-to-end fashion with running polypropylene suture (**Figure 1**). The venous cannulas were then removed, air was evacuated from the mediastinum and pleural spaces, and the sternum and skin incisions were closed. The lambs were extubated and allowed to recover as previously described ^32–34^.

**Figure 1:**
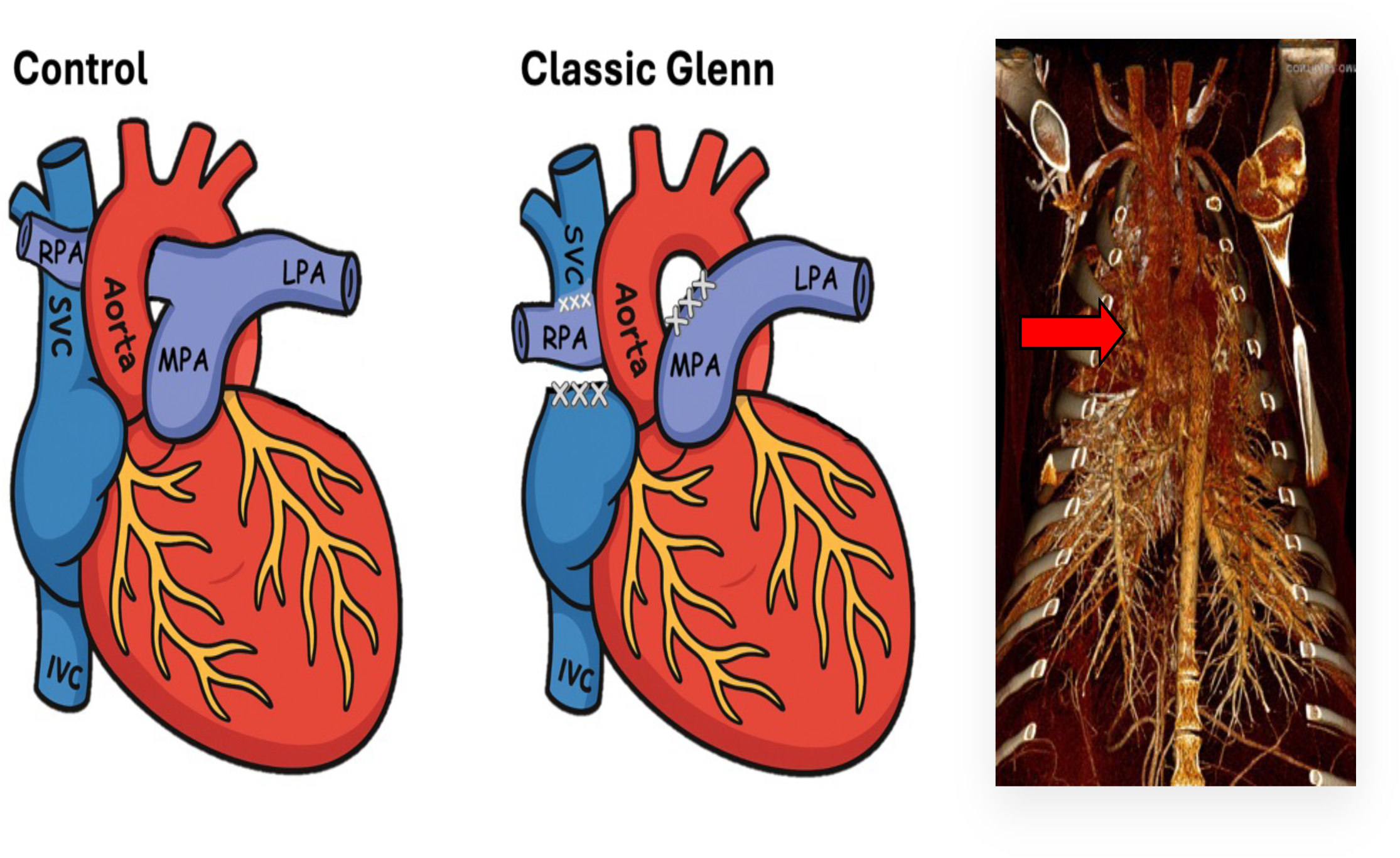
Diagram of normal cardiac anatomy (left); classic superior cavalpulmonary connection (Glenn procedure) (middle); and a representative chest computed tomography angiography (CTA) (right) of our ovine classic Glenn procedure. The superior vena cava (SVC) is connected to the right pulmonary artery (RPA) (**red arrow**), and the RPA is disconnected from the main pulmonary trunk (MPA). Thus, blood flow to the right lung is from the SVC (passive and **non-pulsatile**). LPA=left pulmonary artery.

Hemodynamic measurements were performed eight weeks following the Glenn surgery (n=7). An additional seven age/weight-matched control lambs were studied. At the time of hemodynamic study, lambs were anesthetized, and catheters were placed into the right and left atrium and main pulmonary artery and an ultrasonic flow probe (Transonics Sytems, Ithaca, NY) was placed around the left pulmonary artery to measure pulmonary blood flow, as previously described^28,35^. In all Glenn lambs an additional catheter was placed in the SVC to measure the pressure in the Glenn circulation. In three Glenn lambs, a flow probe was also placed on the right pulmonary artery to measure right (Glenn) blood flow.

At the end of the protocol, all lambs were euthanized with a lethal injection of sodium pentobarbital (150 mg/kg) followed by bilateral thoracotomy as described in the National Institutes of Health Guidelines for the Care and Use of Laboratory Animals. All protocols and procedures were approved by the Committees on Animal Research at the University of California, San Francisco (San Francisco, CA) and the University of California, Davis (Davis, CA).

### Isolated Pulmonary Artery Studies

Third-fourth-generation PA (inner diameters of ∼2-3 mm) were dissected, isolated from the right lung of Glenn and control lambs, and cut into rings as described previously^36^. A continuous recording of isometric force generation was obtained by tying each vessel ring to a force-displacement transducer (Model UC2; Statham Instruments, Hato Rey, PR) that was connected to a recorder (Gould Instrument Systems, Valley View, OH). After the arterial rings were mounted, they were allowed to equilibrate for 20 min in the bathing solution. A micrometer was used to stretch the tissues repeatedly in small increments over the following 45 min until resting tone remained stable at a passive tension of 0.8g as is standard to test vasomotor activity in isolated vessels. Isolated PA were pretreated with indomethacin (10^−5^ M) to prevent the formation of vasoactive prostaglandins and propranolol (10^−6^ M) to block β-adrenergic receptors. To examine the effect of the endothelium-dependent vasodilator acetylcholine (Ach), PAs were first pre-constricted with an EC80 (the concentration required to achieve 80% of maximum constriction) of norepinephrine (NE). Once the response to NE had reached a steady level, cumulative concentration-response curves to Ach (0.001 μM to 100 μM) were obtained by increasing the bath concentration of these drugs in successive steps: the next concentration was added only when the response to the prior concentration had reached a plateau. Vessel rings were used for one experimental protocol and then discarded.

### mRNA extraction and quantitative real-time PCR

Total RNA was isolated from frozen lung tissue samples using the RNeasy mini kit (Qiagen, Netherlands) according to the manufacturer’s protocol. RNA concentration was quantified using a NanoDrop spectrophotometer (ND-1000, ThermoFisher Scientific, Waltham, MA) and reverse transcription was performed with the RNA to cDNA EcoDry Premix (Oligo dT) (Takara Bio, Japan) using 1 μg of total RNA. Quantitative real-time PCR amplification was done in triplicate 20 μL reactions using PerfeCTa SYBR Green SuperMix ROX (Quantabio, Beverly, MA) on an ABI 7900HT Real-Time PCR System (ThermoFisher). Gene-specific primers were designed using the public PrimerQuest Tool software (Integrated DNA Technologies, Skokie, IL). qPCR results were calculated using the comparative C_T_ method as previously described (22) with beta-2-microglobulin as the reference gene and normalized to compare relative changes in mRNA expression in Glenn conditions to control.

### Western blot

Proteins were extracted, and sample concentrations were determined using a SmartSpec 300 spectrophotometer (Biod-Rad, Hercules, CA). Western blot was performed as described previously^37^. 20 μg of protein were loaded per sample and separated by 10% SDS-PAGE. Proteins were subsequently transferred to a polyvinylidene difluoride (PVDF) membrane (Millipore Sigma, Burlington, MA) and blocked with 5% nonfat dried milk in 130mM NaCl and 25mM Tris (TBS, pH 7.5) for 1 hr at room temperature. Blots were incubated overnight at 4°C with primary antibodies against eNOS (BD Transduction Laboratories, Milpitas, CA, USA, 610297), preproET-1 (Invitrogen, Waltham, MA, USA, MA3-005), ECE-1 (Abcam, Cambridge, UK, ab71829), ETRa (Abcam, ab178454) and ETRb (Invitrogen, PA3-066) receptors, and β-actin (Abcam, ab8227) which served as a loading control. This was followed by several washes in TBST and incubation with the species-appropriate horseradish peroxidase or IRDye-conjugated secondary antibody for 1 hour at 4°C. After final washes in TBST, protein bands were visualized by chemiluminescence (SuperSignal West Pico Chemiluminescent Substrate kit, ThermoFisher). Relative protein expression was calculated by band densitometry using the public domain Java image-processing program ImageJ (NIH Image) and normalized to compare relative changes in protein expression in Glenn conditions to control.

### NOx Determinations

To quantify bioavailable NO, NO and its metabolites were determined in lamb lung tissue. In solution, NO reacts with molecular oxygen to form nitrite, and with oxyhemoglobin and superoxide anion to form nitrate. Nitrite and nitrate are reduced using vanadium (III) and hydrochloric acid at 90°C. NO is purged from the solution resulting in a peak of NO for subsequent detection by chemiluminescence (NOA 280; Sievers Instruments, Boulder, CO), as we have previously described^37^. The sensitivity is 1 × 10^−12^ mol, with a concentration range of 1 × 10^−9^ to 1 × 10^−3^ molar of nitrate.

### NOS Activity

NOS activity was measured with the Nitric Oxide Synthase (NOS) Assay Kit (Sigma-Aldrich, St. Louis, MO). Tissue samples were homogenized in PBS and centrifuged at 10,000 x g. The resulting supernatant was mixed with a working reagent that catalyzes the NOS reaction. NO production was measured following the reduction of nitrate to nitrite using the Griess Method. The linear detection range is 0.25 - 25 U/L.

### ET-1 Determinations

Proteins were extracted from lung tissue by homogenization in radioimmunoprecipitation (RIPA) buffer (150mM NaCl, 1% Nonidet P-40, 0.25% sodium deoxycholate, 1mM EDTA, 50mM Tris-HCl) containing a protease inhibitor cocktail (Sigma Aldrich, St. Louis, MO) followed by sonification and centrifugation to collect liquid lysate. Protein concentrations were then determined for lung homogenates via a Bicinchoninic Acid (BCA) Protein Assay Kit (Sigma Aldrich) and for whole cell lysates via Quick Start Bradford Protein Assay (Bio-Rad, Hercules, CA). Endothelin-1 ELISA was performed for both tissue and plasma using a commercial kit according to the manufacturer’s instructions (Enzo Life Sciences, Farmingdale, NY), as we have previously described^38^. The sensitivity is 0.41 pg/mL with a range of 0.78-100 pg/mL. Standard and sample concentrations were calculated by fitting the data to a four-parameter logistic regression.

### Cell Culture

Pulmonary Artery Endothelial Cells (PAECs) were isolated and cultured as previously described^28^. Briefly, primary PAECs were isolated via the explant technique from the right pulmonary artery. A segment of the right PA was placed in a sterile dish containing DMEM with appropriate supplementation. The segment was stripped of adventitia with sterile forceps and opened longitudinally, and the endothelial layer was removed by gentle rubbing with a cell scraper. Cells were grown in culture in appropriate media. After several days, moderate-sized aggregates of endothelial cells were transferred using a micropipette, grown to confluence, and then maintained in culture.

### Cell Proliferation

The procedure for quantifying cell proliferation was performed as previously described^28^. PAECs primarily derived from control and Glenn lambs were treated with 0.25% trypsin solution, washed with PBS, and then counted using a coulter-based cell counter (Moxi Z, Orflo). Next, 6000 PAECs were seeded into 24-well cell culture plates with standard cell culture media. Three distinct cell lines from each experimental group were plated in replicates of 5 per line for each experimental growth time point. At sequential 24-h time points after seeding, cells from each line were trypsinized and counted (as per above) out to 120 hours. This procedure was repeated in triplicate.

### Apoptosis Detection

Cells from the control and Glenn cell lines were seeded into a well plate (∼ 5,000 cells/well) and allowed to grow for 48 hours. The cells were treated with an apoptosis inducer and then fixed and dried. Formamide (a denaturing reagent that denatures DNA only in apoptotic cells) was added and allowed to incubate for 45 minutes. The formamide was removed and the blocking solution was added. After 1 hour, the blocking solution was removed and an antibody mixture, containing antibodies that bind to single stranded DNA was added for 30 minutes. After washing and addition of peroxidase substrate, the cells were read in an ELISA plate reader at 405 nm (ApoStrand ELISA Apoptosis Detection Kit (Enzo Life Sciences, Plymouth Meeting, PA). A higher absorbance at 405 nm indicates a larger percentage of apoptotic cells.

### Angiogenesis Quantification

Matrigel assays were performed in 24-well plates previously coated with 50 µL/cm^2^ of Corning Matrigel Basement Membrane Matrix (Corning #354234; 9 mg/mL, Tewksbury, MA, USA). Cells were seeded (40,000 cells/well, passage 2–5) and adhered for 4–6 h. Four pictures/well were taken using the Cell Imaging System EVOS (Thermo Fisher Scientific, Waltham, MA) and automatic quantification of tube formation was performed with the Image J Angiogenesis Analyzer. Briefly, the sum of nodes and isolated segments were calculated to quantify the capacity of PAECs for tube formation.

### Computed Tomography Imaging

Seven weeks after the Glenn procedure, lambs were placed under a 12-hour enforced fast and were induced and maintained under general anesthesia using IACUC-approved institutional anesthetic protocols. The lambs were placed in dorsoventral recumbency and all CT images were acquired under an enforced 10cm/H20 breath hold to reduce respiratory motion. All CT images were acquired at 120kV and 150mA with a 0.6mm slice thickness (GE Lightspeed Multi-Slice Helical Scanner General Electric Co., Milwaukee, WI). Pre-contrast images were obtained from the thoracic inlet through the entire thorax and included all lung fields. Post-contrast images were obtained similarly fashion after intravenous non-ionic iodinated contrast material (Isovue 370, Bracco Diagnostics Inc., Princeton, NJ) was administered at 2.4mls/kg through a cephalic venous catheter via pressure injector (Medrad Stellant, Bayer HealthCare LLC, Whippany, NJ) at 4ml/second using a 5-second pre-scan delay. Images were viewed in standard and thorax algorithms in 0.6mm slice thickness and all image reconstruction was performed on a dedicated Picture Archiving and Communication System (PACS) workstation (Agfa-Gevaert Group, Greenville, SC).

### Histopathologic Analysis

Dual immunofluorescence for Von Willebrand Factor (VWF, 1:100 dilution; A0082, Dako) and Ki67 (1:100 dilution; M7240, Dako) was carried out on formalin-fixed, paraffin-embedded 5-µm sections from a representative lung sample from four controls and four Glenn. Following citrate pH 6.0 antigen retrieval and serum block, antibodies were incubated overnight at room temperature. VWF was developed with donkey anti-rabbit Alexa Fluor 488 and Ki67 with donkey anti-mouse cyanine (Cy3) (both 1:500 Jackson ImmunoResearch). Coverslips were mounted using Vectashield fluorescent mounting medium with DAPI (Vector Laboratories). Images were visualized and captured with a digital camera mounted on a Nikon Eclipse 80i microscope using NIS-Elements Advanced Research Software v4.13 (Nikon Instruments Inc., Melville, NY). A manual count of all arterial VWF-positive endothelial cells expressing Ki67 on a single 5-µm section was tabulated from images captured at 40x with documentation of the number of pulmonary arteries in the section.

### Human studies

Children with SVHD, living with Glenn physiology for a minimum of 2 years, presenting to the UCSF Benioff Children’s Hospitals for an elective Fontan procedure were included. Systemic arterial blood was obtained from an arterial line placed just prior to their procedure. Children with biventricular heart disease and normal pulmonary blood flow and pressure throughout their life serviced as age-matched controls. Whole blood was centrifuged at 4°C at 3,000xg for 15 minutes. Plasma was isolated and stored at -80° until assayed for ET-1 as described above. The protocol and procedures were approved by the UCSF institutional review board (IRB) and the guardians of all subjects provided written informed consent.

### Statistical Analysis

Differences between treatment groups were compared by the unpaired t-test for normally distributed continuous variables and by Fisher’s exact test for categorical variables. Repeated measures were analyzed by mixed measure ANOVA, with post-hoc comparisons made at discrete timepoints using unpaired t-tests with Bonferroni correction. Data was tested for normality and non-parametric testing was utilized when appropriate. Linear regression analyses were performed to investigate associations between human plasma ET-1 levels and demographic and hemodynamic parameters. Significant outliers, defined as data points greater than two standard deviations from the mean were excluded from the analysis. A p<0.05 was considered significant. Statistical Analysis was done using R software^39^.

## Results

Eight weeks after the Glenn procedure, lambs (Glenn and age/weight-matched controls) underwent a terminal hemodynamic study (**Table 1**). There were no differences in age, sex, weight, hemoglobin, or oxygen saturation between the groups. Mean pulmonary arterial pressure was higher in Glenn lambs but remained within normal limits (p<0.05). Glenn pressure was low (6.4±1.7 mmHg), and as expected, left lung blood flow was higher, and right lung blood flow was lower in Glenn lambs compared to controls (p<0.05). All other baseline hemodynamics were similar between Glenn and control lambs.

**Table 1:** Baseline Demographics and Hemodynamics.

|  | **CONTROLS (N=7)** | **GLENN (N=7)** |
| --- | --- | --- |
| **Age (days)** | 105.1 ± 8.9 | 119.0 ± 21.8 |
| **Weight (kg)** | 35.0 ± 5.3 | 29.7 ± 7.9 |
| **Hemoglobin (gm/dl)** | 12.0 ± 0.6 | 11.0 ± 1.1 |
| **Oxygen Saturation** | 95.0 ± 1.6 | 92.6 ± 2.8 |
| **HR (beats/min)** | 163.9 ± 42.1 | 161.3±27.2 |
| **MSAP (mmHg)** | 105.1 ± 18.5 | 98.2±14.0 |
| **MPAP (mmHg)** | 12.5±2.4 | 16.7±3.1* |
| **LAP (mmHg)** | 5.3 ± 2.9 | 4.0±1.3 |
| **RAP (mmHg)** | 3.7 ± 2.2 | 2.9±2.3 |
| **Glenn (SVC) Pressure (mmHg)** | NA | 6.4±1.7 |
| **Left PA Flow (L/min/kg)** | 0.040±0.014 | 0.064±0.014* |
| **Right PA Flow (L/min/kg) (n=3)** | 0.058±0.021† | 0.012±0.002* |

HR= heart rate; MSAP=mean systemic arterial pressure; MPAP=mean pulmonary artery pressure; LAP=left atrial pressure; RAP=right atrial pressure; SVC=superior vena cava; PA=pulmonary artery. In Glenn lambs, MPAP reflects the pressure in the left lung. \* P<0.05 vs. control.
† Calculated assuming a 55:45 right to left lung pulmonary blood flow ratio.

### Isolated Pulmonary Artery Studies

Relaxations to the endothelium-dependent vasodilator, Ach, were used to test for responsiveness to endogenously produced NO (a functional indicator of EC function). As seen in **Figure 2**, isolated PAs from the right lung of Glenn lambs demonstrated impaired relaxation to Ach compared to PAs isolated from the right lung of control lambs (p <0.05).

**Figure 2:**
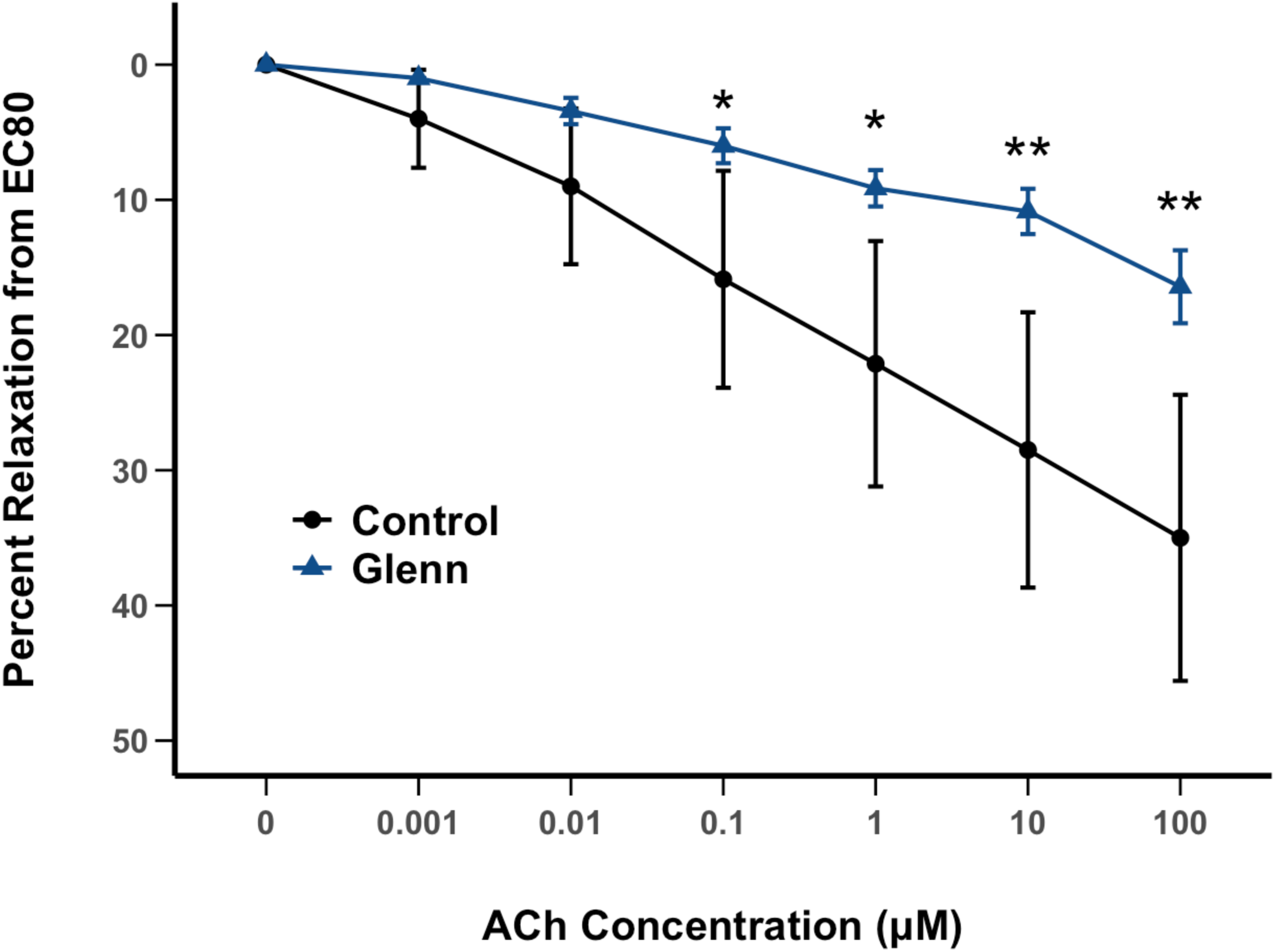
Isolated vessel pulmonary arterial (PA, 3^rd^-4^th^ generation) reactivity to the endothelium-dependent vasodilator acetylcholine (Ach) following norepinephrine pre-constriction from control and Glenn right lungs (n=7 lambs per group, average of two vessels per lamb). Downward deflection notes relaxation. Values are mean±SD. *p<0.05; **p<0.01.

### NO Signaling

As seen in **Figure 3 A, B**, right lung Glenn tissue eNOS mRNA and protein levels were decreased compared to the right lung of controls (p<0.05). eNOS activity (0.34±0.01 vs. 0.35±0.05 U/L normalized to eNOS protein; control vs. Glenn, n=4) was similar between the groups. Lung tissue NO metabolites (NOx), an indirect determinant of bioavailable NO, were decreased in the right lung of Glenn lambs compared to the right lung of controls (p<0.05) (**Figure 3C**).

**Figure 3:**
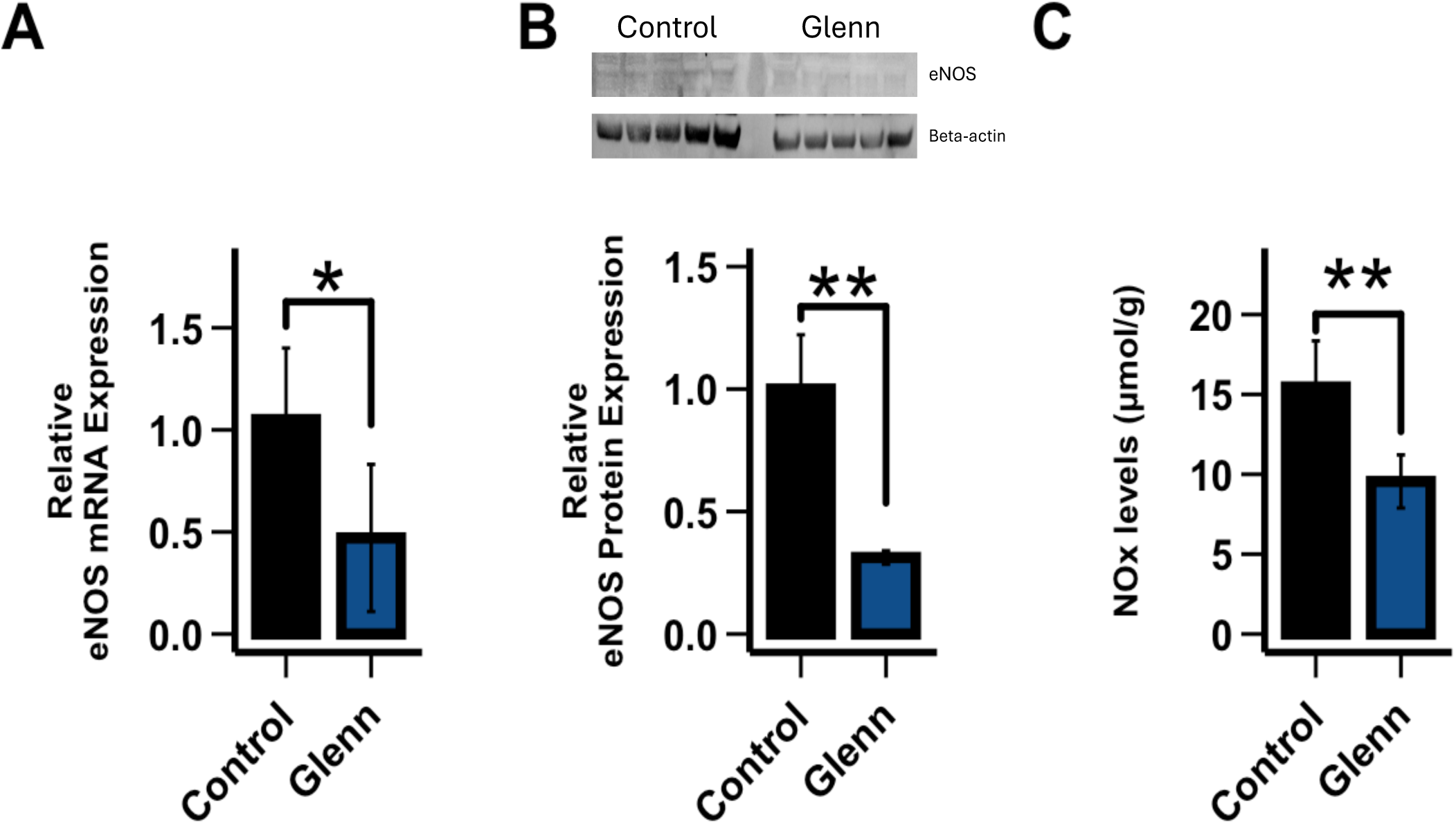
Compared to the right lung of age-matched control lungs, quantitative real-time PCR (qPCR) on right lung Glenn tissue demonstrates that eNOS RNA expression is decreased in right Glenn lungs (**A**)(n=5 per group). Western blot analysis demonstrates that right lung tissue eNOS protein levels are decreased (**B)** (n=5 per group). This correlates with decreased right lung tissue levels of NO metabolites in Glenn lambs compared to the right lung of controls (**C**) (n=6 per group, NOx via chemiluminescence). Values are relative to control values designated as 1. Values are mean ± SD. *p<0.05, **p<0.01).

### ET-1 Signaling

As seen in **Figure 4 A, B**, plasma and right lung tissue ET-1 levels were increased in Glenn lambs compared to controls (n=6 per group, p<0.05). Right lung Glenn tissue Prepro-ET-1 mRNA and protein levels were increased compared to the right lung of controls (**Figure 4 C, D)** (p<0.05). Endothelin Converting Enzyme (ECE-1) lung tissue protein levels were increased in Glenn lambs (**Figure 4 E)** (p<0.05), while ETa and ETb receptor protein expressions were similar between the groups (**Figure 4 F, G**).

**Figure 4:**
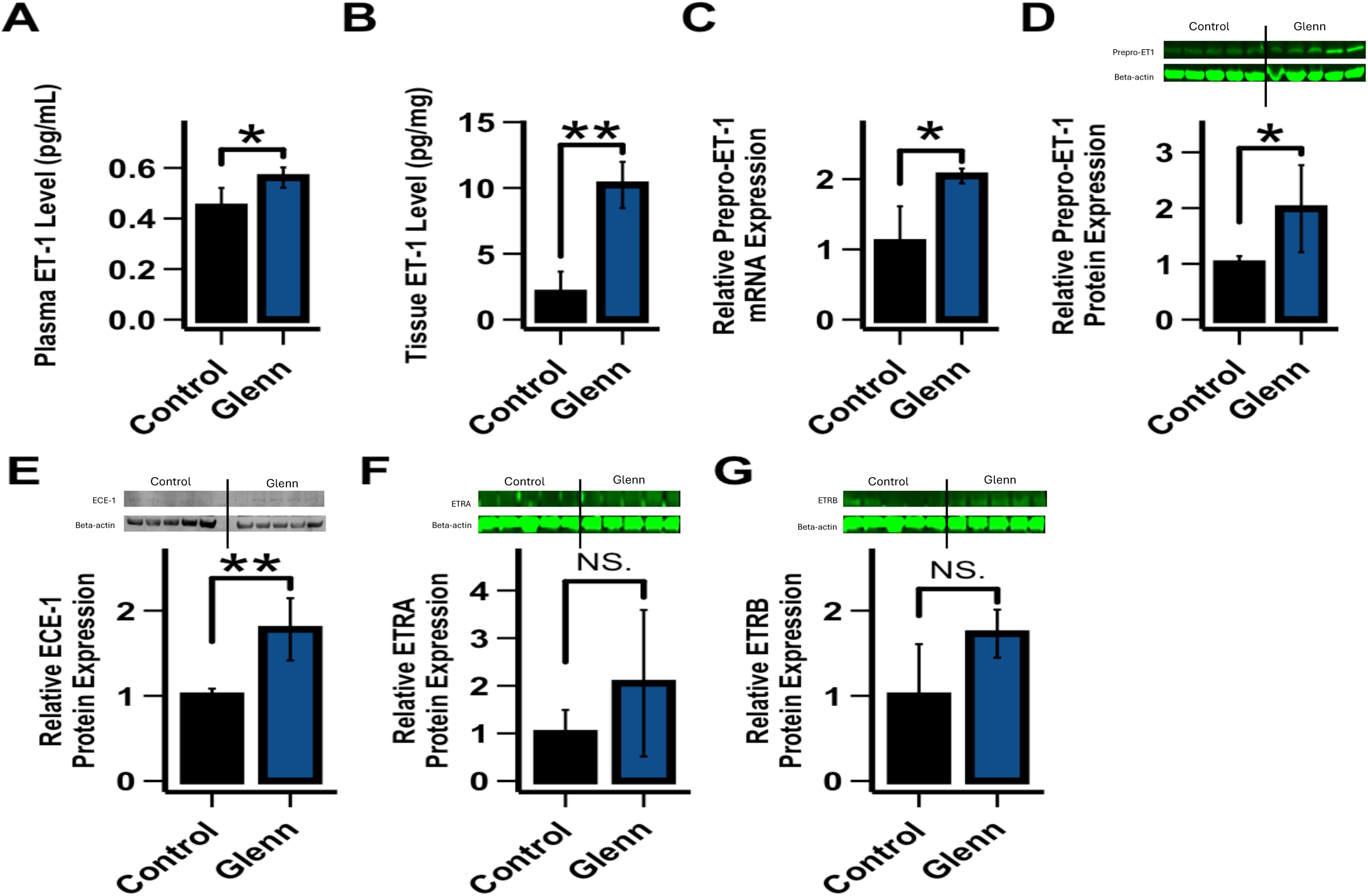
Both plasma (**A**) and right lung tissue (**B**) ET-1 levels are increased in Glenn lambs (n=6 per group). Compared to the right lung of age-matched control lungs, quantitative real-time PCR (qPCR) on right lung tissue demonstrates that Prepro-ET-1 RNA expression is increased in right Glenn lungs (**C**)(n=5 per group). Western blot analysis demonstrates that right lung tissue Prepro-ET-1 protein levels are increased (**D)** (n=5 per group). ECE-1 protein levels (**E**) were increased in the right Glenn lung compared to controls, while ETRA (**F**) and ETRB (**G**) protein levels were similar between groups. qPCR and protein values are relative to control values designated as 1. Values are mean ± SD. *p<0.05, **p<0.01.

### PAEC Functional Characteristics

We next defined the functional characteristics of PAECs derived from the right lung of Glenn lambs and compared them to the right lung of age-match control lambs. To quantify PAEC proliferation, we performed cell counting of cultured cells over a 5-day period. We found that PAECs derived from Glenn lambs (**Figure 5A**) demonstrated significantly increased proliferation compared to control PAECs (p<0.05). Histopathologic analysis confirmed increased EC proliferation in the vessels of Glenn lungs (**Figure 6**). To quantify apoptosis, we utilized an ELISA Apoptosis Detection Kit. Glenn PAECs had a lower percentage of apoptotic cells than controls (p<0.05), indicating apoptotic resistance in the right lung of Glenn lambs. (**Figure 5B**). To characterize angiogenesis, we used a tube formation assay in growth factor–restricted Matrigel. PAECs from Glenn lambs exhibited a significant increase in angiogenesis than control animals after 72 hours in Matrigel, as quantified by the number of branch points (p<0.05)(**Figure 5C,D**). Consistent with other models of PVD^40^, these data collectively demonstrate that PAECs derived from the right lung of Glenn lambs are pro-proliferative, anti-apoptotic, and pro-angiogenic compared to PAECs derived from control lambs.

**Figure 5:**
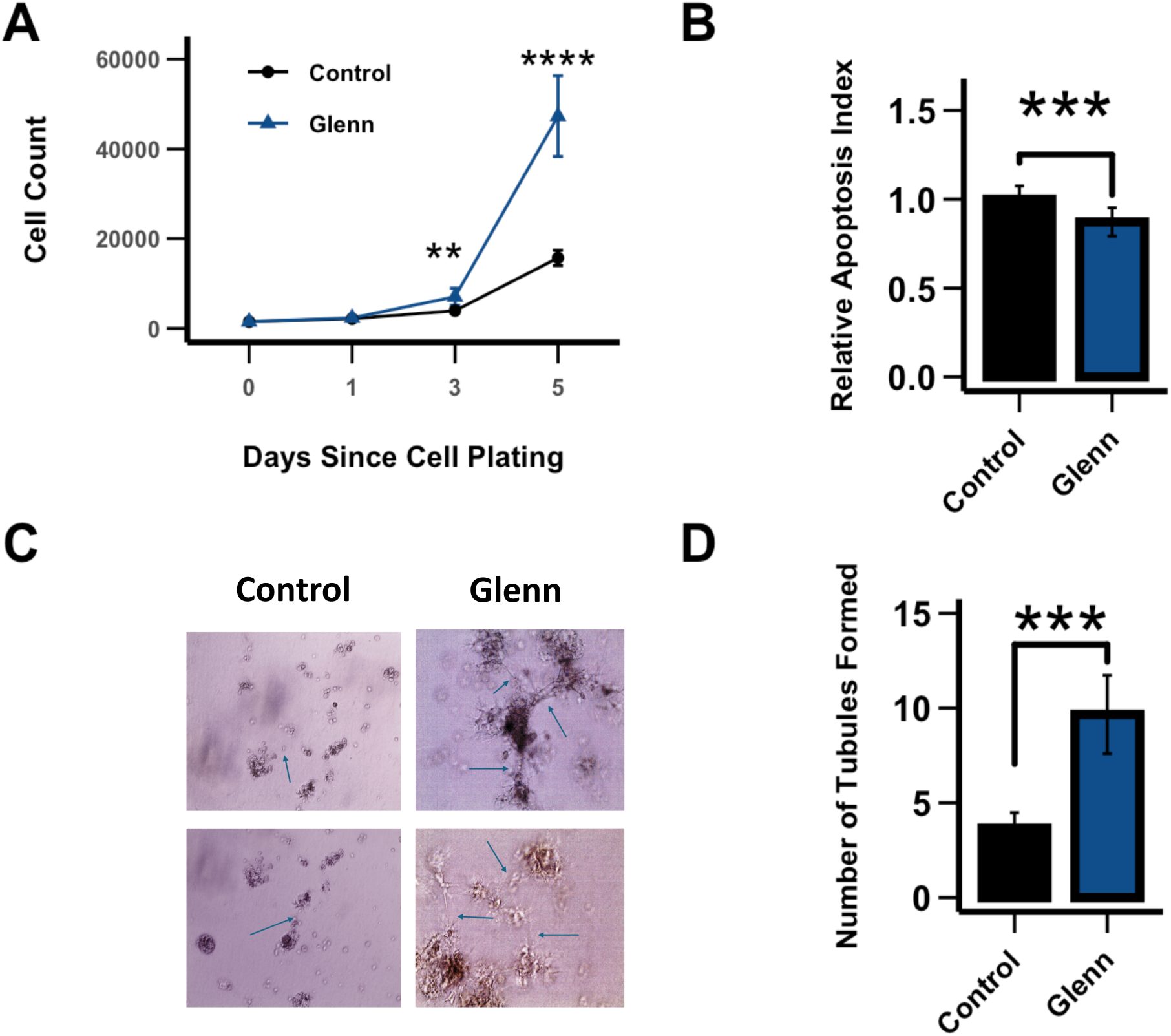
Pulmonary artery endothelial cell (PAEC) functional characteristics: cell proliferation, apoptosis and angiogenesis. (**A**) PAEC cell proliferation assay was performed by cell counting over a 5-day period (n=3 per group). (**B**) Apoptosis tendency was quantified using the ApoStrand ELISA Apoptosis Detection Kit. PAECs from the right lung of Glenn lambs had a lower relative apoptosis index (0.87±0.08) than did PAECs from the right lung of control lambs (1.0±0.08). (**C, D**) PAECs’ angiogenic capacity was assessed using Matrigel assay as quantified by the number of branch points at 72 hrs. PAECs from Glenn right lungs had a greater rate of angiogenesis (9.7±2.0) than the right lung of control lungs (3.7±0.8). (**B**) Representative images, arrows designate branch points; (**D**) cumulative values. Values are mean±SD. n = 3 lambs per group run in triplicate. **p<0.01, ***p<0.001, ****p<0.0001.

**Figure 6:**
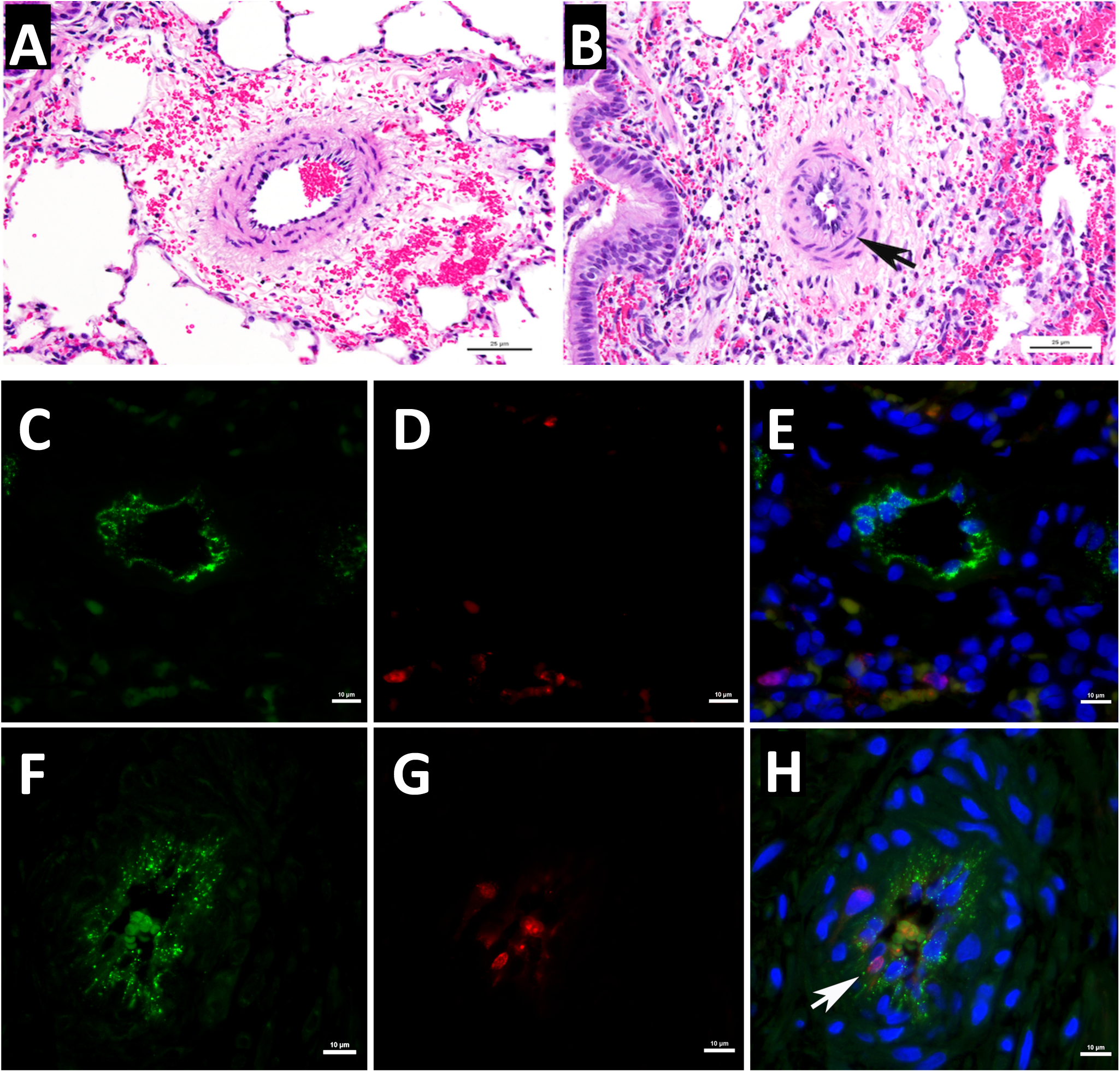
Compared to pulmonary arteries within control lungs (**A**) there is prominence of the endothelium within pulmonary arteries of the Glenn lungs (**B, arrow**), which also shows wall infiltration by individual inflammatory cells (hematoxylin and eosin; n=4 per group). Dual immunofluorescence for Von Willebrand Factor (green) and Ki67 (red) demonstrates increased proliferating cells within the pulmonary artery endothelium (arrow) of Glenn lungs (**F-H**) compared to age-matched controls (**C-E**). Nuclei counterstained with DAPI (blue).

### Human Studies

Plasma ET-1 concentrations were determined in seven children with SVHD, who had previously undergone a Glenn procedure. These children had Glenn physiology (non-pulsatile pulmonary blood flow) for > 3.5 years at the time of sampling, just prior to undergoing a Fontan procedure. Comparisons were made with seven age-matched control patients who had bi-ventricular congenital heart disease with normal pulmonary blood flow (pulsatile) and normal pulmonary hemodynamics. There were no differences in age between the two groups (4.9 ± 0.9 years-Glenn vs. 6.8 ± 5.0 years- control). As seen in **Figure 7**, compared to age-matched controls, circulating plasma ET-1 levels were more than double in children with Glenn physiology (7.1 ± 1.8 pg/mL vs 3.0 ± 0.6 pg/mL, p<0.001). There was no correlation between the ET-1 level and age, weight, or baseline hemodynamics by linear regression analysis.

**Figure 7:**
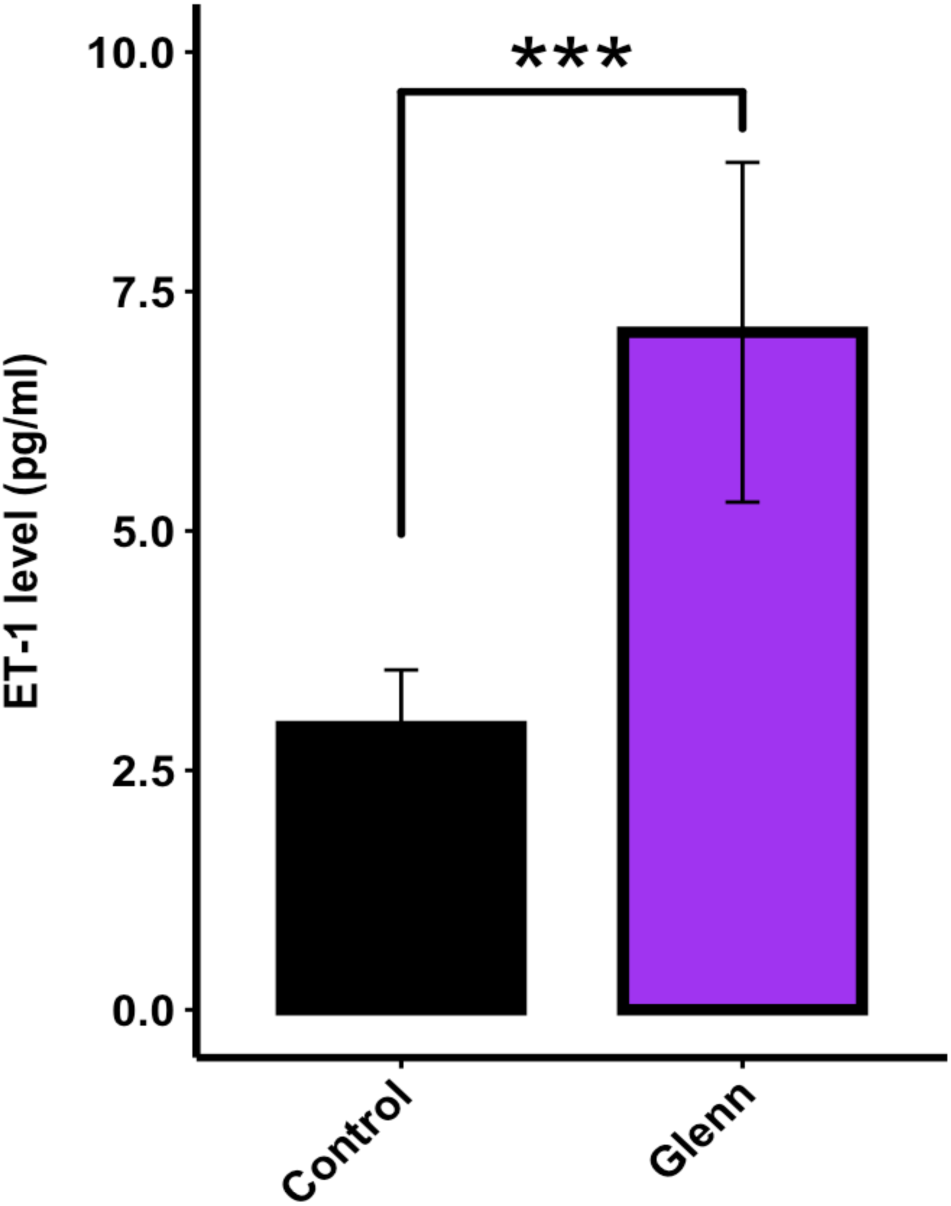
Plasma ET-1 levels (ELISA) **<u>in children with SVHD</u>** who were all s/p Glenn procedures before undergoing a Fontan procedure, and age-matched control with bi-ventricular congenital heart disease. All Glenn children had Glenn physiology (non-pulsatile, passive pulmonary blood flow) for a minimum of 3.5 years. Values are mean±SD. N=7 Glenn; N=7 age-matched bi-ventricular controls. *** p<0.001.

## Discussion

Morbidities associated with pulmonary vascular disease in SVHD, including upper extremity swelling, hypoxemia, and decreased cardiac output are well appreciated^16,17,20,21,41^. However, the preceding pulmonary vascular EC dysfunction contributing to these morbidities has not been well characterized, and its timing and mechanism are not well appreciated. Lopes et al. demonstrated altered levels of endothelial markers in patients with SVHD years following a Glenn procedure^24^. This included increased plasma levels of von Willebrand factor antigen and tissue-type plasminogen activator factor, and decreased thrombomodulin compared to controls. Importantly, all 10 of these patients were deemed inappropriate to proceed to a Fontan due to pulmonary artery abnormalities and/or increased pulmonary artery pressure^24^. In addition, Natarajan et al. demonstrated elevated ET-1 plasma levels in 18 children ∼2 years following their Glenn procedure, just prior to their Fontan^25^. Interestingly, compared to age-matched controls, Frank et al. demonstrated decreased plasma ET-1 levels in 55 patients with SVHD just prior to their Glenn procedure^23^. In the present study, we utilized an ovine model of a classic Glenn to reveal early and extensive evidence of pulmonary vascular EC dysfunction, which includes a marked attenuation of endothelium-dependent relaxation of isolated pulmonary arteries, decreased lung tissue bioavailable NO, and increased ET-1. In addition, PAECs derived from these vessels demonstrated a hyperproliferative, angiogenic, anti-apoptotic phenotype classically associated with pulmonary vascular disease^40^. Importantly, histopathologic analysis confirmed EC hyperproliferation in the vessels of Glenn lambs. Notably, this degree of EC dysfunction was present only eight weeks following the Glenn procedure. Thus, despite the appreciation that pulmonary vascular disease occurs in patients with the Fontan circulation following years of hypoxemia, polycythemia, inflammation, and non-pulsatile pulmonary blood flow, these data suggest that pathology is extensive just weeks following the Glenn procedure. Importantly, these aberrations are occurring in the setting of non-pulsatile pulmonary blood flow in the absence of hypoxemia or polycythemia, suggesting that non-pulsatile blood flow is paramount to this pathology.

Endothelial cells (EC) are constantly under the influence of hemodynamic forces including: (1) shear stress, the tangential friction force acting on the vessel wall due to blood flow; (2) hydrostatic pressure, the perpendicular force acting on the vascular wall; and (3) cyclic strain, the circumferential stretch of the vessel wall. Mechanosensors on EC detect these forces and transduce them into biochemical signals that trigger vascular responses. The regulation of endothelial gene expression through biomechanical forces is a critical determinant of normal and abnormal vascular tone, remodeling, and development^26,42,43^. For example, the principal physiological stimulus for NO synthase (NOS) under normal conditions is laminar shear stress^44^. However, chronic alterations in mechanical forces associated with vascular remodeling disrupt these normal responses and result in endothelial dysfunction, including decreased bioavailable NO and altered vascular tone^36,37^. The Glenn procedure initiates a low-flow, non-pulsatile source of pulmonary blood flow^9^. Lack of pulsatility has been previously recognized as a potential driver of EC dysfunction. For example, non-pulsatile left ventricular mechanical support devices are associated with systemic EC dysfunction, while pulsatile devices preserve EC function and microcirculatory blood flow^45,46^. Utilizing a classic Glenn model in pigs, Henaine et al. demonstrated impaired endothelium-dependent pulmonary relaxation and decreased eNOS expression in the right lung, three months following the procedure. However, these aberrations were absent in a group of pigs that had a secondary source of pulsatile pulmonary blood flow to the right lung^47^. Importantly, in humans that underwent a Glenn procedure with an additional source of pulsatile pulmonary blood flow, Kurotobi et al. demonstrated that an impaired pulmonary vasodilating response to the endothelium-dependent vasodilator acetylcholine negatively correlated with the pulmonary pulse pressure, suggesting that the lack of pulsatility was responsible for the pulmonary vascular EC dysfunction^48^. In addition, a recent meta-analysis of studies comparing the impact of maintaining a source of pulsatile pulmonary blood flow during the Glenn procedure suggested improved pulmonary artery development compared to those children without a source of pulsatile blood flow^49^ . The current study supports the hypothesis that lack of pulsatile pulmonary blood flow following surgical palliation for SVHD is a major driver of pulmonary vascular EC dysfunction and subsequent pulmonary vascular disease. However, the potential additive/independent effects of chronic hypoxia, inflammation, and/or polycythemia warrant further investigations.

Hallmarks of endothelial dysfunction include decreases in the vasodilating and smooth muscle cell anti-proliferative molecule NO and increases in the vasoconstricting and pro-proliferative peptide ET-1. In the pulmonary vasculature of Glenn lambs, the decrease in NO signaling was manifest as an impairment in Ach (which requires the EC to make NO to induce dilation) mediated pulmonary relaxation in isolated pulmonary arteries, decreased eNOS mRNA and protein levels in lung tissue, and decreased lung tissue NO metabolites, an indirect marker of bioavailable NO (**Figures 2 and 3**). In addition, lung tissue eNOS activity was unchanged compared to control lambs suggesting a lack of post-translational modifications. However, it is well appreciated that eNOS can be post-translationally modified in manners that affect its function by a variety of conditions^50^. Although our data suggest that the decrease in NO production is likely related to transcriptional decreases as opposed to post-translational modifications, it is noteworthy that the NOS activity assay is performed at Vmax, potentially masking post-translational modifications of activity including co-factor availability. The active ET-1 is a 21-amino acid polypeptide that arises from a larger polypeptide, prepro-ET-1, that is cleaved by the endothelin converting enzyme (ECE-1)^51^. In the pulmonary vasculature of Glenn lambs, the increase in ET signaling was manifest as increased lung tissue and plasma ET-1 levels, an increase in prepro-ET1 mRNA and protein lung tissue expression and an increase in ECE-1 tissue protein levels (**Figure 4**). These data suggest that both the transcriptional increase in prepro-ET-1 and its converting enzyme participate in the demonstrated increases in ET-1 levels. In addition, both tissue ETa and ETb receptor expression were unchanged. This contrasts with a study of patients who died following a failed Fontan, in which lung ET receptor expression was increased in autopsy specimens^52^. However potential changes in receptor binding affinity and location cannot be excluded. Importantly, the increase in ET-1 levels was confirmed in plasma levels of children with Glenn physiology (**Figure 7**).

Regulation of angiogenesis and EC proliferation in the pulmonary vasculature is complex, with both adaptive homeostatic features, in addition to maladaptive phenotypes that contribute to the development of pulmonary vascular disease. For example, highly proliferative, anti-apoptotic endothelial cells are present in advanced vascular remodeling that include plexiform lesions and complex intravascular lesions with luminal obstruction^53–55^. PAECs from the Glenn right lung demonstrated a hyper-proliferative, anti-apoptotic phenotype (**Figure 5**). However, the histologic examination of lungs from Glenn lambs revealed moderate medial hypertrophy and EC proliferation, with no evidence of either endothelial obliteration of the vessel lumen or plexiform lesions. Interestingly, in a separate ovine model of early pulmonary vascular disease secondary to increased pulmonary blood flow and pressure, we saw a similar PAEC hyperproliferative, anti-apoptotic phenotype without advanced vascular remodeling^28,35^. Thus, the hyperproliferative, anti-apoptotic phenotype seen in these models may be precursor to more advanced pathology. Similarly in our previous model of CHD with increased pulmonary blood flow and pressure, PAECs demonstrated a pro-angiogenic phenotype^28^. In that setting, angiogenesis could have represented an adaptive response to incorporate the increased flow, since these lambs developed increased vessel density compared to controls^35^. In the current Glenn model, the angiogenic response may be independent of altered blood flow patterns. It is well recognized that patients with Glenn physiology, and this ovine Glenn model, develop pulmonary arterial-venous malformations (PAVMS)^32,34,56^. Importantly, we confirmed PAVMs in the right lung of all the Glenn lambs in this study by agitated saline echocardiography (data not shown). Clinical observations suggest that PAVM development is related to the lack of first-pass hepatic venous return to the lungs during Glenn physiology, suggesting that a hepatic-derived inhibitor of angiogenesis is pivotal to the proangiogenic environment^57^. Thus, the PAEC pro-angiogenic phenotype demonstrated in our current Glenn model may reflect this pathology.

Limitations of the current investigation are noteworthy. For example, although a significant strength of the study is that the large animal model attempts to isolate the effect of non-pulsatile pulmonary blood flow on EC function, clinically patients with SVHD often also suffer from hypoxemia and elevated hematocrits. However, the potential additive and/or synergistic effects of hypoxemia and polycythemia are excluded from the current study and warrant further investigations. Similarly, the potential role of aberrant flow-induced inflammation warrants consideration^58^ . In addition, the potential effect of the differential source of pulmonary blood flow (i.e. SVC only, thereby lacking hepatic venous return), a likely significant contributor of EC pathology, is not evaluated and requires further study. It is noteworthy that the source of our PAECs was from proximal pulmonary arteries since these are most exposed to the mechanical forces associated with differential blood flow patterns. However, PAECs from small resistance vessels are also critical, and require further study. Lastly, the potential effect that the pulmonary vascular EC dysfunction induced by the Glenn physiology might have on distant vascular beds is a crucial question that also warrants further investigation.

## Conclusion

In summary, utilizing an in-vivo large animal model of Glenn physiology, coupled with ex-vivo and in-vitro investigations, we demonstrate early and extensive evidence of pulmonary vascular EC dysfunction associated with the non-pulsatile pulmonary blood flow initiated with Glenn physiology. In the absence of hypoxemia and polycythemia in our in-vivo model, these data strongly suggest that the low, non-pulsatile flow associated with Glenn physiology plays a pivotal role in the development of EC dysfunction. Further investigation into the aberrations in mechanosensing associated with these flow abnormalities, and their downstream mechanisms that induce EC dysfunction may reveal important novel therapeutic targets. The use of endothelial-based pulmonary vasodilator therapies has been reported in SVHD. However, reports have been predominantly in adult patients, years following their final staged surgery (i.e. Fontan). Although the safety of pulmonary vasodilator treatment in this population has been consistent, efficacy in small series has been inconsistent^31,59–65^. Importantly, two large randomized controlled trials in adults with SVHD following a Fontan did not meet their primary endpoints. For example, there was no difference in exercise capacity in a 52-week phase 3 trial of Macitentan, an endothelin receptor antagonist, and no difference in oxygen consumption at peak exercise in a 26-week phase 3 trial of Udenafil, a phosphodiesterase type 5 inhibitor^66,67^. However, these studies represent treatment strategies in adults following years of exposure to non-pulsatile flow-induced EC dysfunction. Clinical trials focused on the prevention of EC dysfunction initiated at the onset of non-pulsatile aberrant flow patterns warrant future consideration.

## Acknowledgements

The authors would like to thank Rachel Hutchings, Christian Vento, Shannon Cheung, Reza Kheirkhahi Oqani, and Kelsea Lee for their expert technical assistance.

This work was supported by National Institutes of Health (NIH) grant T32HL160508 (Hyde, Smith, Hwang), K12HD105250 (Smith), K08HL148512 (Boehme), R01HL133034, R01 HD072455 (Maltepe), P01HL146369 (Black, Fineman, Maltepe, Wang), and Additional Ventures Grants P0583323 (Hwang) and P0583322 (Datar).

## Conflict of Interest

No conflicts of interest, financial or otherwise, are declared by the authors.

All animal protocols and procedures were approved by the Committees on Animal Research at University of California, San Francisco and Davis, and comply with the National Institutes of Health Guidelines for the Care and Use of Laboratory Animals. All human protocols and procedures were approved by the Committee on Animal Research at University of California, San Francisco (San Francisco, CA). Appropriate consent was obtained.

All data will be made available upon request by the corresponding author.

